# Arginase 1 is negatively regulated by β-catenin signaling in the lung

**DOI:** 10.1101/137539

**Authors:** Joseph A. Sennello, Annette S. Flozak, Alexander V. Misharin, Cara J. Gottardi, Anna P. Lam

## Abstract

We previously demonstrated that mice lacking the Wnt co-receptor, Lrp5, had attenuated pulmonary fibrosis in the bleomycin model. We found that Arginase 1 (*Arg1)*, an enzyme that converts L-arginine to urea and ornithine, was markedly elevated in *Lrp5*^-/-^ lungs compared with wild-type mice after bleomycin injury. We show that this induction is not apparently due to the expression of Th2 cytokines, IL-4 and IL-13, but instead is due to Wnt/β-catenin signaling, which negatively regulates Arg1 expression in lung macrophages. Although Arg1 expression in macrophages has been used to define an alternatively activated phenotype, flow cytometry analysis of alveolar and interstitial macrophage sub-populations in *Lrp5*^-/-^ lungs 14 days after bleomycin injury revealed no clear evidence of skewing from a classical to an alternatively activated phenotype. Upregulation of *Arg1* expression and arginase activity might diminish lung arginine levels with consequent alterations in collagen or cytokine production. However, dietary supplementation of bleomycin-treated *Lrp5*^-/-^ mice with the Arg1 substrate, L-arginine, failed to alter lung collagen content or cytokine levels 21 days after bleomycin injury. These findings demonstrate that Arg1 is negatively regulated by β-catenin signaling in macrophages, raising the possibility that Wnt signaling directs alterations in immune cell metabolism that may be relevant to lung repair after injury.

## INTRODUCTION

Arginase 1 (Arg1) is an enzyme that converts L-arginine to urea and ornithine, and is best known for its function in the liver, where it catalyzes the final step of the urea cycle required for ammonia detoxification [1, 2]. Ornithine can be used to generate polyamines, glutamate, and proline, from which collagen is synthesized [3-9]. Outside the liver, Arg1 is inducibly expressed by macrophages, where it competes with iNOS to control production of nitric oxide in classically activated macrophages stimulated by Th1 cytokines. Its expression can be robustly stimulated by the Th2 cytokines, IL-4 and IL-13, and a Th2 immune response has been shown to be associated with accumulation of Arg1-expressing macrophages [9-11] that contribute to the development of fibrosis [4, 10, 12-18].

In Idiopathic Pulmonary Fibrosis (IPF), developmental pathways, such as Wnt/β-catenin, have been implicated in the cycle of aberrant wound repair, resulting in persistent fibrosis [19]. The Wnt/β-catenin pathway is best known in determining cell fate, such that disruption of Wnt/β-catenin signaling results in early embryonic demise, axis malformation, and aberrant lung development [20-22]. In the adult tissue, Wnt/β-catenin signaling plays critical roles in hematopoietic stem cell maintenance, gut epithelial cell self-renewal [23, 24], and bone density [25-29]. In the liver, Wnt/β-catenin signaling supports hepatocyte proliferation and ammonia detoxification, whereby a gradient of β-catenin signaling separates glutamine synthesis and urea formation [30, 31]. Together, these studies support the paradigm in which Wnt/β-catenin signaling is exquisitely cell-type and cell-context dependent.

Work from our lab has demonstrated that global loss of the Wnt co-receptor Lrp5 protects from bleomycin-induced pulmonary fibrosis [32]. We identified *Arg1* as one of the top upregulated genes in these bleomycin injured *Lrp5*^-/-^ lungs, and investigated whether this increased *Arg1* expression reflected altered lung metabolism of L-arginine and a strong Th2 cytokine milieu, or its negative regulation by β-catenin signaling in lung macrophages.

## RESULTS

We previously demonstrated that mice lacking Lrp5 globally were protected from bleomycin-induced pulmonary fibrosis [32] and that microarray analysis of these lungs at day 14 after administration of bleomycin revealed enrichment for gene pathways related to the immune response [33]. Of the genes with increased expression, we confirmed that *Arg1* was highly upregulated by real-time PCR analysis in a separate cohort of mice (Figure 1A). *Arg1* mRNA expression was highly increased in *Lrp5*^-/-^ compared to *Lrp5*^+/+^ bleomycin-injured lungs at day 14 (17.1±6.2 fold versus 1.2±0.4 fold). This increased *Arg1* expression likely also reflected increased activity of arginase-1, as urea formation was increased in *Lrp5*^-/-^ compared to *Lrp5*^+/+^ lungs at day 14 (3.65±0.58 versus 2.55±0.39 mg/ml; Figure 1B). In uninjured saline-treated mice, expression of *Arg1* in *Lrp5*^-/-^ lungs was also increased compared to *Lrp5*^+/+^ littermates (7.33±9.21 fold versus 0.97±0.21; Figure 1C). Together, these data suggested that global knockout of the Wnt co-receptor Lrp5, which results in a diminished capacity to transduce canonical Wnt signals [32], is associated with increased *Arg1* expression in the lung. While β-catenin signaling was known to inhibit *Arg1 expression* in the liver [30, 31], this relationship was not yet established in the lung.

**Figure 1.**
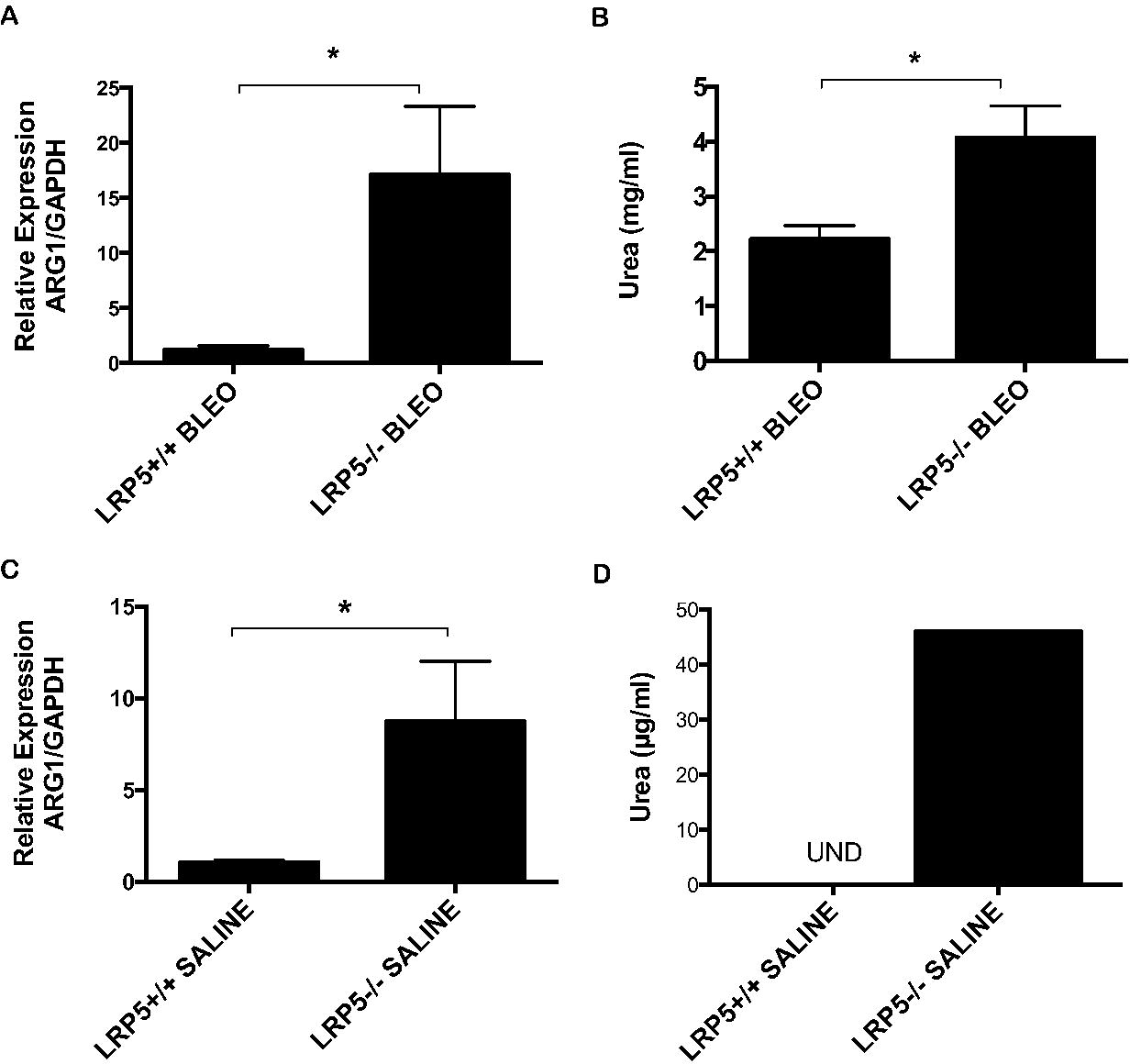
*Arg1* gene expression and activity are increased in mouse lungs with global loss of the Wnt co-receptor Lrp5. (**A**) *Lrp5*^-/-^ and *Lrp5*^+/+^ mice were treated with intratracheal bleomycin, and lungs were harvested on day 14 for RNA extraction. *Arg1* gene expression was measured by real time qPCR and normalized to *GAPDH* (n=5-9 animals per condition, \**p*<0.05). (**B**) *Lrp5*^-/-^ and *Lrp5*^+/+^ lungs harvested on day 14 after bleomycin injury were analyzed for arginase activity assay by measuring urea (n=4 animals per condition, \**p*<0.05). Saline-treated *Lrp5*^-/-^ lung tissue manifests greater (**C**) *Arg1* gene expression and (**D**) Arg1 protein activity, assessed by urea assay, at baseline than *Lrp5*^+/+^ lungs.

Arg1 expression can be stimulated by the Th2 cytokines and the resultant production of polyamines can subsequently affect expression of pro-inflammatory factors [8, 9]. Therefore, we assessed whether the increased Arg1 expression was due to an increase in basal Th1 or Th2 cytokine levels, which were measured from whole lung tissue by ELISA. Saline treated *Lrp5*^-/-^ mice demonstrated decreased levels of IL-4 (201±17 pg/ml versus 322±30 pg/ml at day 6, p=0.02; 214±29 pg/ml versus 317±50 pg/ml at day 14, p=0.17), but there were no differences in basal IL-13, TNF-α, and IFN-γ levels compared to *Lrp5*^+/+^ littermates (Figure 2). These results suggest that increased *Arg1* expression due to global loss of Lrp5 was not due to increased basal Th2 immune response in the uninjured lung. Next, we examined whether bleomycin-induced lung injury in *Lrp5*^-/-^ mice affected production of the Th2 cytokines and resultantly increased Arg1 expression. We found no differences in levels of IL-4 or IL-13 from *Lrp5*^-/-^ compared to *Lrp5*^+/+^ lungs on days 6 and 14 after bleomycin injury. However, TNF-α and IFN-γ levels were increased in bleomycin treated *Lrp5*^-/-^ lungs compared to *Lrp5*^+/+^ controls at day 6 (177±16 versus 118±7 pg/ml and 91±5 versus 71±4 pg/ml, respectively; Figure 2), but these differences dissipated by day 14 after bleomycin treatment. These findings indicate that global loss of Lrp5 resulting in increased *Arg1* expression after bleomycin injury is not associated with an increased Th2 inflammatory response.

**Figure 2.**
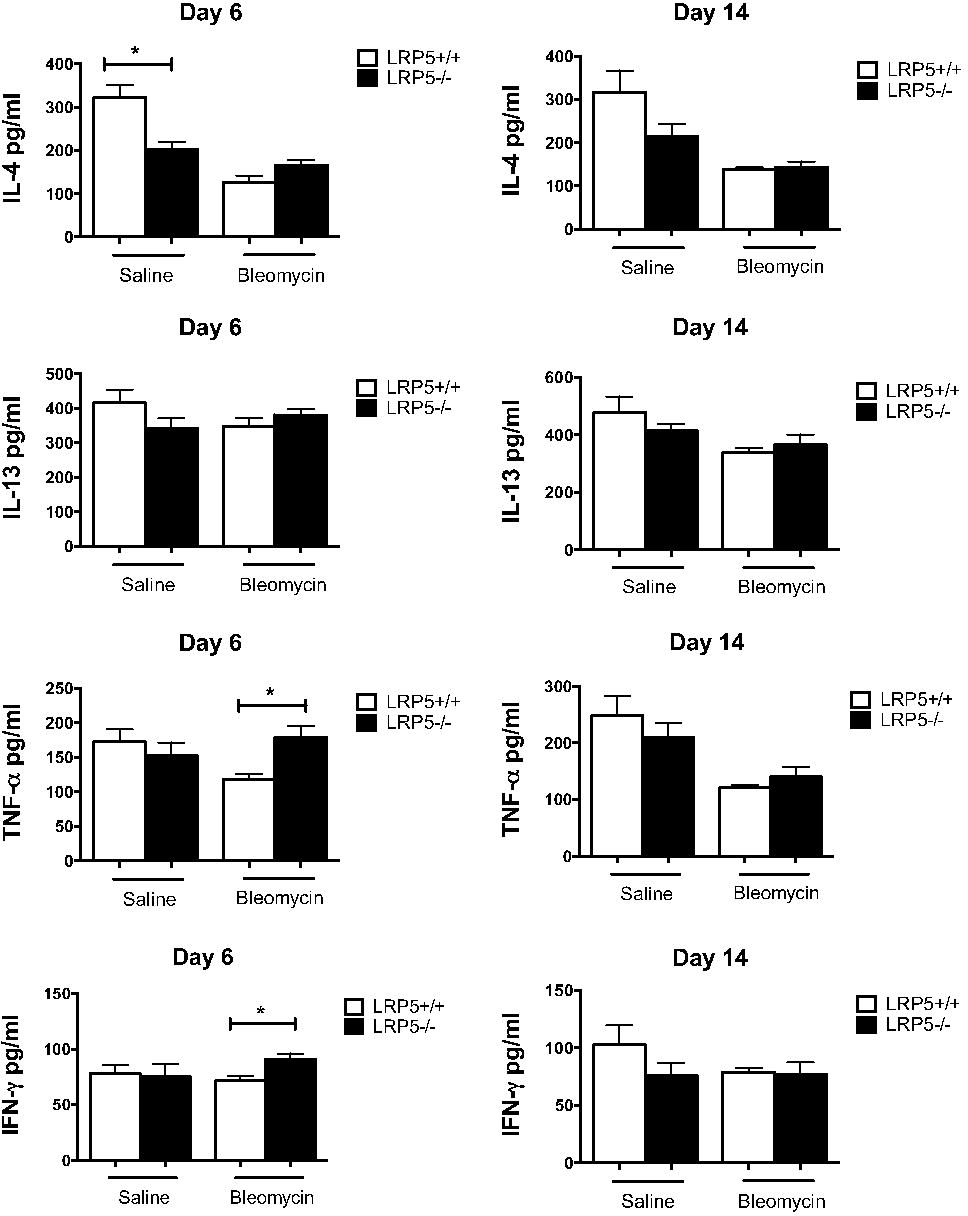
Basal IL-4 level is decreased in lungs lacking Lrp5, but only TNF-α and IFN-γ are increased transiently in *Lrp5*^-/-^ lungs early after bleomycin injury. *Lrp5*^-/-^ and *Lrp5*^+/+^ lungs were homogenized on day 6 or day 14 after intracheal bleomycin or saline treatment. The Th2 cytokines, IL-4 and IL-13, and the Th1 cytokines, TNF-α and IFN-γ, were quantified by ELISA (n=3-4 animals per condition, \**p*<0.05).

Outside the liver, *Arg1* is expressed in macrophages [6, 7, 34], where it has been reported to be a marker of an alternatively activated macrophage phenotype [9, 11, 35, 36]. To determine whether the increase in Arg1 observed 14 days post-bleomycin injury was due to an increase in the number of alternatively activated macrophages in the lungs, we analyzed macrophage differentiation by flow cytometry using standard markers for myeloid cell activation. We found that alveolar macrophages (CD11c^+^ CD11b^-^ CD64^+^ Siglec F^+^) from bleomycin-treated *Lrp5*^-/-^ lungs demonstrated decreased expression of CD80, a marker for classical activation, compared to *Lrp5*^+/+^ lungs (5396±370 versus 7695±470 MFI; Figure 3), but no differences in expression of Relma and CD206, markers of alternative activation. In contrast, *Lrp5*^-/-^ interstitial macrophages (CD11b^+^ MHC II^+^ CD64^+^ CD24^-^ Siglec F^-^) analyzed day 14 after bleomycin injury demonstrated increased expression of both the classical activation marker CD40 (847±79 versus 536±26 MFI) and the marker of alternative activation, RELMa (342±20 versus 233±23 MFI). Expression of CD80, CD86, and CD206 were also increased in *Lrp5*^-/-^ interstitial macrophages but did not reach statistical significance (Figure 3). Thus, while increased Arg1 expression is associated with a mixed activation phenotype in *Lrp5*^-/-^ interstitial macrophages, *Lrp5*^-/-^ alveolar macrophages demonstrated neither classical nor alternative activation, indicating that increased Arg1 expression did not strongly reflect alternative activation of macrophages in the lungs.

**Figure 3.**
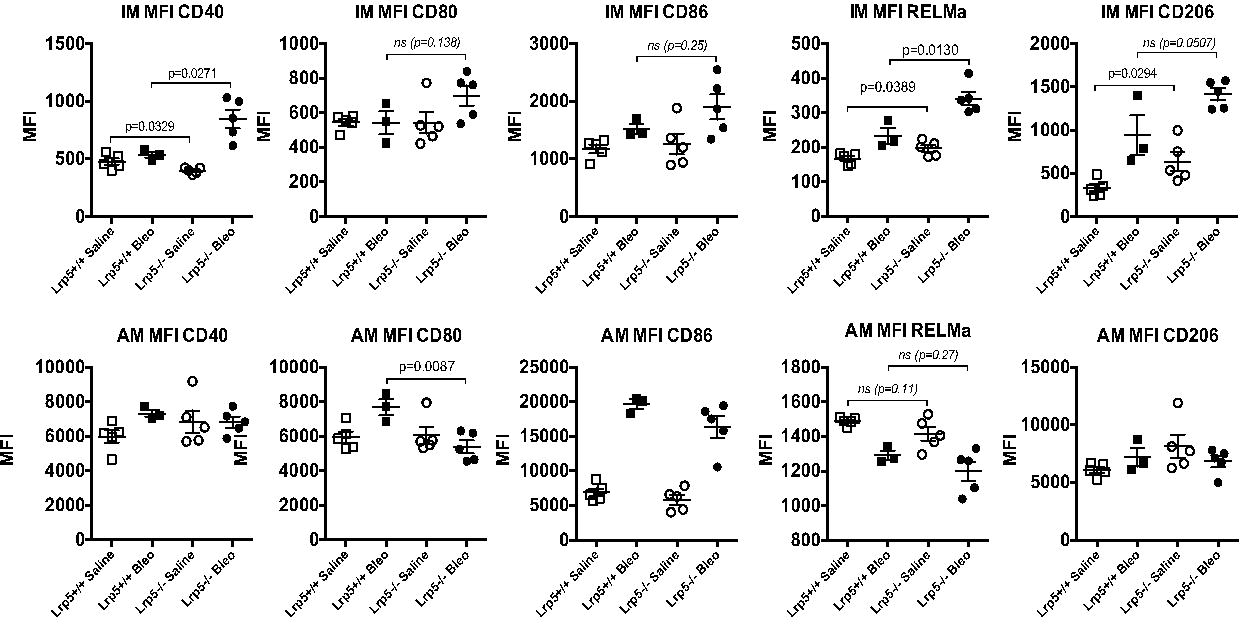
Interstitial macrophages but not alveolar macrophages from *Lrp5*^-/-^ lungs exhibit mixed activation phenotype 14 days after bleomycin injury. *Lrp5*^+/+^ and *Lrp5*^-/-^ mice were treated with bleomycin or saline, and whole lungs were harvested on day 14 after bleomycin for isolation of lung leukocytes. Flow cytometry analysis was performed on interstitial macrophages (IM) and alveolar macrophages (AM) to examine expression of CD40, CD80, CD86, RELMa, and CD206. Results are expressed as mean fluorescent intensity (MFI) (n=5 animals per condition).

We reasoned that increased Arg1 expression and arginase activity after bleomycin injury might deplete local levels of the arginase substrate, L-arginine, in the lungs. Given the reported roles of arginase and L-arginine in inflammation and wound healing [9], we investigated whether supplementation of L-arginine in *Lrp5*^-/-^ mice with elevated *Arg1* gene expression would reverse the observed protection and exacerbate the development of bleomycin-induced pulmonary fibrosis. As expected, we observed that L-arginine supplementation increased arginase activity and urea formation in *Lrp5*^+/+^ lungs (2.63±0.45 versus 1.31±0.05 mg/ml, p=0.04). However, L-arginine supplementation decreased urea formation in *Lrp5*^-/-^ mice, in which arginase activity was already elevated (3.75±0.95 versus 1.71±0.33 mg/ml, p=0.06; Figure 4). In bleomycin-injured *Lrp5*^-/-^ mice, L-arginine supplementation did not increase the soluble collagen content in the lung or bronchoalveolar lavage fluid total protein levels at day 21 (Figure 4). Furthermore, L-arginine supplementation resulted in only modest but not statistically significant elevation of both Th1 and Th2 cytokines at day 21: TNF-α (76.5±22.7 versus 50.9±9.4 pg/ml, p=0.2) and IFN-γ (92.8±10.4 versus 60.1±21.4 pg/ml, p=0.2), as well as IL-4 (112.4±22.6 versus 74.4±20.7 pg/ml, p=0.2) and IL-13 (266.8±46.7 versus 203.3±47.2 pg/ml, p=0.3) (Figure 5). Thus, while L-arginine replacement in Arg1-expressing cells can alter endpoints relevant to fibrogenesis *in vitro* [9, 37], our findings demonstrate that dietary L-arginine supplementation does not significantly impact the development of lung fibrosis or cytokine production after bleomycin injury in mice.

**Figure 4.**
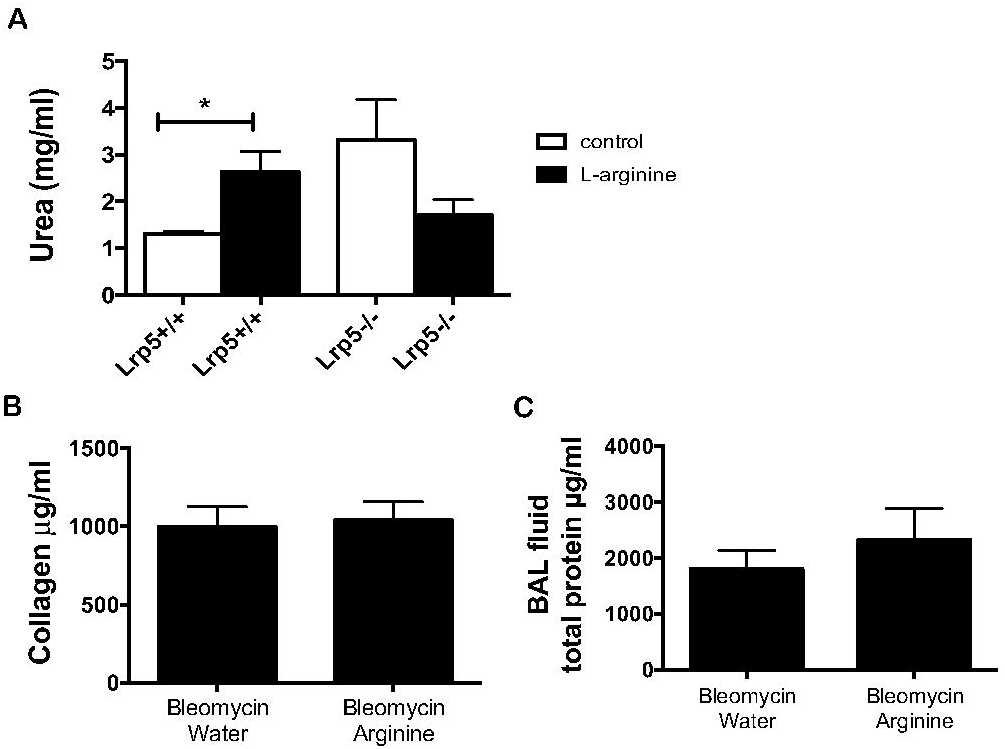
Supplemental L-arginine does not alter lung fibrosis in *Lrp5*^-/-^ mice after bleomycin injury. L-arginine or sucrose was added to the drinking water of *Lrp5*^-/-^ mice for 14 days prior to administration of intratracheal bleomycin. *Lrp5*^+/+^ and *Lrp5*^-/-^ mice supplemented with L-arginine or sucrose were harvested on day 21 after bleomycin administration and analyzed for arginase activity assay by measuring (**A**) urea and for bleomycin-induced injury by measurement of (**B**) total lung collagen content by hydroxyproline assay and (**C**) total protein in BAL fluid (n=3-5 animals per condition, \**p*<0.05).

**Figure 5.**
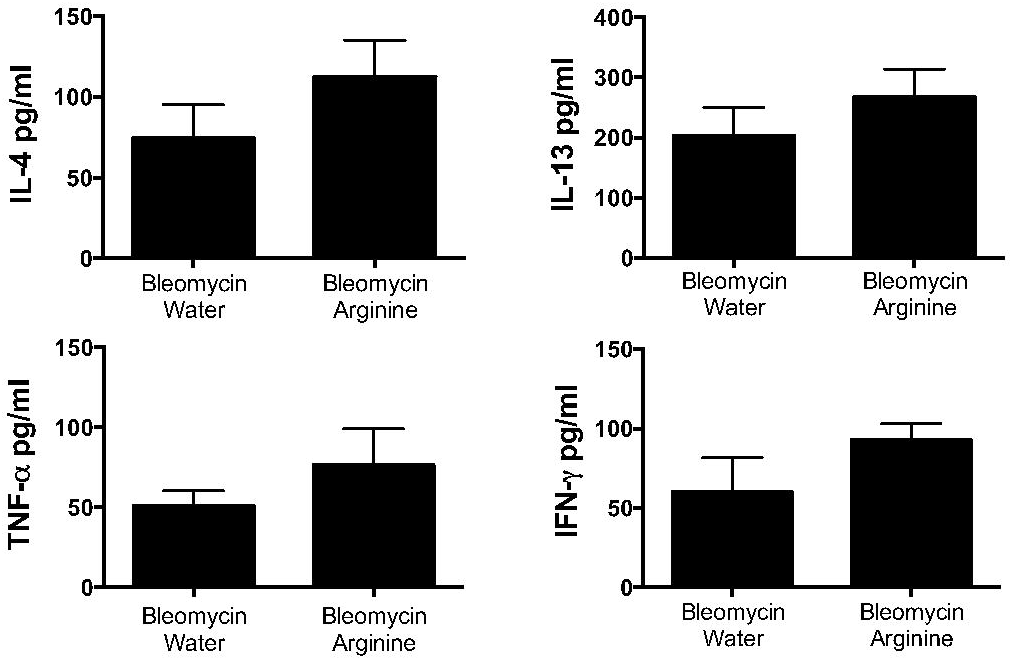
L-arginine supplementation does not significantly alter Th1 or Th2 cytokine production in *Lrp5*^-/-^ lungs after bleomycin injury. *Lrp5*^-/-^ mice supplemented with L-arginine or sucrose in their drinking water for 14 days prior to administration of intratracheal bleomycin. Levels of the cytokines IL-4, IL-13, TNF-α, and IFN-γ were measured by ELISA using homogenized lung lysates harvested on day 21 after bleomycin (n=3-5 animals per condition).

To determine whether the increased Arg1 expression in *Lrp5*^-/-^ mice was due to negative regulation by Wnt/β-catenin signaling, we sought to investigate the effect of inhibiting β-catenin signaling in the MH-S lung macrophage cell line [38, 39]. We confirmed that MH-S macrophages increased *Arg1* gene expression in response to IL-4 stimulation (288.9±24.5 versus 0.62±0.2-fold; Figure 6A). Inhibition of β-catenin signaling with the compound iCRT5, which prevents β-catenin from binding to its transcriptional cofactors [32, 40], increased *Arg1* gene expression (2.66±0.3 versus 0.62±0.2-fold; Figure 6A), consistent with findings in the *Lrp5*^-/-^ lungs, where diminished Wnt/β-catenin activity resulted in increased *Arg1* gene expression at baseline and after bleomycin injury (Figure 1). IL-4 increased expression of the downstream β-catenin target gene *Axin2*, which was abrogated by iCRT5 (2.98±0.3-fold and 1.35±0.3-fold, respectively; Figure 6B), confirming that iCRT5 specifically inhibits β-catenin signaling in these cells and supports previous data suggesting that IL-4 can activate canonical Wnt signaling [41]. Lastly, genetic loss of β-catenin by Cre-recombinase in CD11c macrophages led to increased basal *Arg1* expression *in vitro* (3.89±1.7 versus 1.00±0.44-fold; Figure 6C). This increase in *Arg1* expression persisted regardless of whether these cells were stimulated with Wnt3a or IL-4 (9.49±3.7 versus 1.89±0.64-fold and 1942.8±693.6 versus 900.1±290.8-fold, respectively; Figure 6C). Together these findings indicate that Wnt/β-catenin signaling negatively regulates *Arg1* gene expression in the lung, likely within a subpopulation of macrophages.

**Figure 6.**
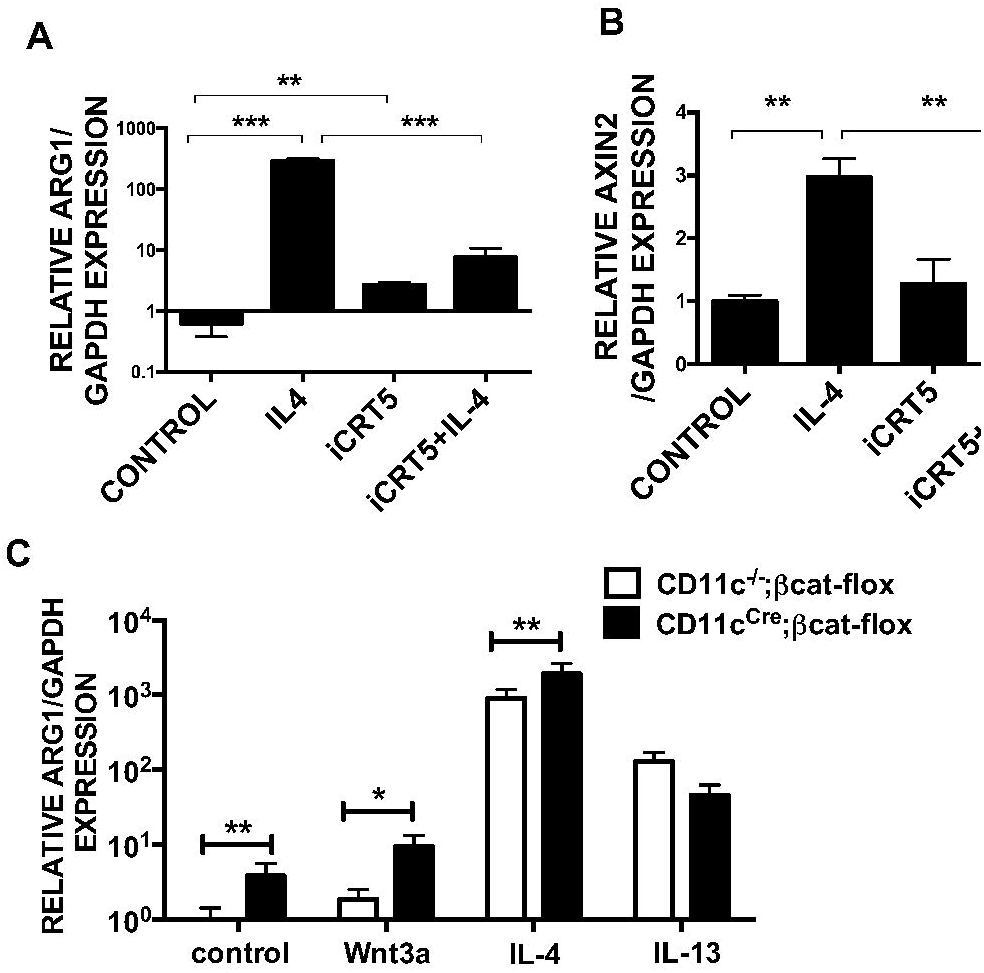
*In vitro* inhibition of β-catenin signaling in macrophages increases *Arg1* gene expression. MH-S macrophages were treated with the small molecule inhibitor of β-catenin activity, iCRT5, and/or IL-4 for 24 hours and then RNA was extracted. Gene expression was determined by real time qPCR analysis for (**A**) *Arg1* and (**B**) *Axin2*, a direct target of β-catenin signaling (n=3 samples per condition, \**p*<0.05, \*\**p*<0.01, \*\*\**p*<0.001). (**C**) Bone marrow cells isolated from *CD11c^Cre^;β-catenin^flox^* mice and *CD11c^-/-^;β-catenin^flox^* littermates underwent *in vitro* differentiation into macrophages. Bone marrow derived macrophages were treated with recombinant Wnt3a, IL-4, or IL-13 for 24 hours prior to RNA isolation. Gene expression of *Arg1* was measured by real-time qPCR and normalized to *GAPDH* (n=3 samples per condition, \**p*<0.05, \*\**p*<0.01, \*\*\**p*<0.001).

## DISCUSSION

Work from our lab previously demonstrated that loss of the Wnt co-receptor, Lrp5, protects from bleomycin-induced pulmonary fibrosis [32]. To determine the mechanism by which diminished β-catenin signaling attenuates the fibrotic response, we sought to identify genes most altered in the lungs of bleomycin-treated *Lrp5*^-/-^ mice relative to *Lrp5*^+/+^ controls. We found that *Arg1* was one of the most highly upregulated targets in *Lrp5*^-/-^ lungs. Arg1 is best known for its role in liver, where it converts L-arginine to urea and ornithine, the former of which facilitates removal of excess nitrogen from the body [1, 2]. Arg1 is also transiently induced in macrophages, where ornithine can be used to generate polyamines, glutamate, and proline, which are useful building blocks for a variety of processes mediated by macrophages [4-6]. Since Arg1 expression can be induced by Th2 cytokines [5, 9], we reasoned that levels of IL-4 and/or IL-13 might be elevated in bleomycin-injured Lrp5^-/-^ mice. We found, however, that IL-4 levels were actually diminished in uninjured *Lrp5*^-/-^ lungs, and neither IL-4 nor IL-13 was altered after bleomycin injury, suggesting a Th2 cytokine-independent mode of regulation. Indeed, inhibition of β-catenin signaling by both chemical (iCRT5) and genetic (targeted removal of β-catenin in CD11c-positive bone marrow derived macrophages by Cre-recombinase) approaches led to elevated *Arg1* expression in macrophages, consistent with studies in liver showing that Arg1 can be negatively regulated by Wnt/β-catenin signaling [30, 31, 42, 43]. Although the means through which Wnt/β-catenin signaling inhibits Arg1 expression is unclear, it is likely indirect given that the core transcriptional components in this pathway serve to promote gene activation [42, 43]. Recent work from Ken Cadigan’s group using flies has identified a number of Wnt/β-catenin-repressed genes that require TCF and a non-canonical HMG-DNA binding consensus site (-WGAWAW-) [44, 45]. However, the extent to which a similar mechanism exists in mammalian systems is not known. Thus, as was observed in liver, we find that Arg1 is also negatively regulated by β-catenin signaling in the lung.

Outside the liver, *Arg1* is induced in macrophages and is a marker of alternative activation, which has been associated with tissue-injury promoting and resolving activities depending on context [37, 46-48]. To assess whether increased Arg1 expression was associated with alternative activation of lung macrophage subpopulations, we performed flow cytometry analysis on *Lrp5*^-/-^ alveolar and interstitial macrophages. Although alterations in macrophage activation were observed, we failed to detect expression of cell surface markers that suggested clear skewing towards an alternative activation phenotype. While these data may be consistent with recent evidence showing that increased alternative activation of lung macrophages leads to worsened fibrosis after bleomycin injury [11, 49], it is important to point out that the simple paradigm of non-overlapping, stable classically and alternatively activated subsets of macrophages is in the process of being replaced by a model that accommodates greater plasticity macrophage phenotypes and cytokine profiles [35, 50]. Studies are underway to determine how β-catenin signaling alters the differentiation state and activities of lung macrophage subpopulations.

Arginase-1 converts L-arginine into L-ornithine, which can be converted to polyamines and proline, the latter of which is required for the production of collagen and the former for cell proliferation/DNA synthesis. This relationship suggested the possibility that elevated Arg1 activity might limit the development of fibrosis through locally depleting L-arginine, a key substrate for the production of proline-rich collagen or the proliferation of cell types that drive fibrosis [51]. To determine whether L-arginine could impact the development of lung fibrosis, we supplemented the diets of bleomycin-injured *Lrp5*^-/-^ mice with L-arginine, to assess whether this would restore wild-type levels of collagen and overall fibrosis. However, supplementation with L-arginine did not increase the levels of collagen in the lungs of *Lrp5*^-/-^ mice, even though L-arginine supplementation was sufficient to alter total lung arginase activity. Thus, in so far as L-arginine supplementation of cell cultures can drive cellular outcomes relevant to fibrosis *in vitro* [9, 37], L-arginine supplementation at the dietary level cannot obviously affect these processes. There were also no significant differences in the levels of Th1 or Th2 cytokines. These findings are consistent with those observed in Tie2-Cre mice sensitized and challenged with ovalbumin, where targeted Arg1 ablation had no effect on expression of IL-4, IL-13, IL-10 or lung content of IL-4, IL-10, TNF-α and IFN-γ [52]. Thus, these data suggest that Arg1 itself may not be a key contributor to the fibrotic phenotype.

While L-arginine is a substrate for arginase, it is also metabolized by inducible nitric oxide synthase (iNOS), producing nitric oxide (NO) and citrulline. Previous work suggested that arginase might control NO synthesis by limiting the intracellular arginine pool, such that inhibition of NO synthesis in macrophages was associated with activation of arginase with reciprocal competition between the activation of iNOS and arginase [5, 6]. Chemical inhibition of NO production using aminoguanidine has been shown to limit bleomycin-induced pulmonary fibrosis [53, 54], while genetic deletion of iNOS abolished pulmonary fibrosis in a chronic ova-albumin model but had no effect on the inflammatory response [55]. While our studies did not measure NO levels directly to determine whether increased Arg1 activity resulted in limited L-arginine available for NO production, the exogenous repletion of L-arginine to a level that was able to alter total lung arginase activity, and presumably provided sufficient substrate for NO production, did not impact the fibrotic response. We acknowledge that there may still be a contribution by the iNOS/NO pathway. However, our collective findings better support a novel role for Wnt/β-catenin signaling in directing alterations in immune cell metabolism for tissue repair, consistent with a recent study by Monticelli et al [51].

In summary, we show that Arg1, a top upregulated gene in the lungs of bleomycin-injured Lrp5^-/-^ mice that show reduced fibrosis [32], is negatively regulated by Wnt/β-catenin signaling in macrophages. Given our recent evidence that β-catenin signaling in CD11c-positive lineages is sufficient to antagonize the resolution of fibrosis [33], we reason that identifying the full complement of Wnt/β-catenin target genes in these cells will shed important light on how macrophage differentiation contributes to lung repair after injury.

## METHODS

### Cell culture and animals

All procedures were approved by the Northwestern University Animal Care and Use Committee. C57B/L6 mice and *CD11c*^*cre*^ mice were obtained from Jackson Laboratory (Bar Harbor, ME); *Lrp5*^-/-^ mice from Bart O. Williams (Van Andel Research Institute, Grand Rapids, MI). Recombinant Wnt3a, IL-4, IL-13, and GM-CSF were obtained from Peprotech (Rocky Hill, NJ); iCRT5 was kindly provided by Ram Dasgupta (NYU, New York, NY).

### L-arginine supplementation and bleomycin administration

Specific pathogen free C57Bl/6-Lrp5 null mice and wild-type littermate controls weighing 25-30 g were provided supplemental L-arginine in drinking water containing 2.5% L-arginine (Sigma Aldrich) with 5.0% sucrose. L-arginine supplementation began 14 days prior to bleomycin administration. Mice were lightly anesthetized using isoflurane and given bleomycin (APP Pharmaceutics-Schaumberg, IL) at 1.5 U/kg body weight or 50μl of 0.9% sterile saline via intra-tracheal catheter. Euthanasia was performed using two methods, consisting of cervical dislocation followed by removal of heart and lungs, after complete anesthesia was ensured with administration of pentobarbital.

### Quantification of mRNA

Total RNA was extracted from tissue using RNeasy Kit (Qiagen, Valencia, CA) according to manufacturer’s instructions. Real time qPCR primer sequences are as follows: Mouse GAPDH forward primer 5’-TTGTGATGGGTGTGAACCACGAGA and reverse primer 5’GAGCCCTTCCACAATGCCAAAGTT; mouse ARG1 forward 5’-AACACGGCAGTGGCTTTAACCTTG and reverse 5’-AAGAACAAGCCCTTGGGAGGAGAA; mouse AXIN2 forward 5’-ACCTCAAGTGCAAACTCTCACCCA and reverse 5’-AGCTGTTTCCGTGGATCTCACACT. Real-time PCR (qPCR) was performed using iQ SYBR Green Supermix (Biorad, Hercules, CA) with the MyiQ Single Color Real-Time PCR Detection System and software (Bio-Rad). GAPDH was used as the internal control. The relative change in gene expression was calculated using the 2^-ΔΔCt^ method [56].

### Arginase activity

Lung arginase activity was measured as previously described [57].

### ELISA

IL-4, IL-13, TNF-α, and IFN-γ immunoassays were performed following the manufacturer’s protocol (R&D Systems, Minneapolis, MN).

### Flow cytometry, cell sorting, isolation of bone marrow derived cells and lung leukocytes

Lung leukocytes were isolated using GentleMACS dissociator (Miltenyi, Auburn, CA) as previously described [58]. Bone marrow derived macrophages were generated from whole bone marrow cells isolated from the femurs and tibias of mice, followed by in vitro differentiation in DMEM supplemented with 10% fetal bovine serum (FBS), 1% penicillin/streptomycin, and 20% L929 supernatant containing mouse granulocyte–macrophage colony-stimulating factor. Flow cytometry and cell sorting was performed using LSR II flow cytometer (BD Biosciences, San Jose, CA) as previously described [58].

### Statistics

Statistical analysis was performed by Student’s *t* test or one-way analysis of variance (ANOVA) followed by Bonferroni’s multiple comparison test. A *p* value of less than 0.05 was considered significant. Statistical calculations were performed using GraphPad Prism software (Graph Pad Software Inc, La Jolla, CA).

## Acknowledgements

We would like to thank Ram Dasgupta for the iCRT5 compound.

